# Actin cytoskeleton dynamics affect replication of Human Metapneumovirus

**DOI:** 10.1101/2022.09.30.510364

**Authors:** Pamela Elizabeth Rodríguez, Pedro Ignacio Gil, Jorge Augusto Cámara, Alicia Cámara, María Gabriela Paglini

**Affiliations:** Laboratory of Influenza and other Respiratory Viruses, Virology Institute “Dr. J. M. Vanella”. Facultad de Ciencias Médicas. Universidad Nacional de Córdoba. Córdoba. Córdoba. Argentina; Laboratory of Cell Biology of Viral Infection, Virology Institute “Dr. J. M. Vanella”. Facultad de Ciencias Médicas. Universidad Nacional de Córdoba. Córdoba. Córdoba. Argentina; Laboratory of Neurophysiology, Instituto de Investigación Médica Mercedes y Martín Ferreyra, INIMEC-CONICET, Universidad Nacional de Córdoba, Córdoba, Argentina

**Keywords:** Virus isolation, Replication cycle, Actin filaments, Cytochalasin D, Respiratory disease

## Abstract

Human Metapneumovirus (hMPV) is responsible for viral respiratory infection with clinical and epidemiological relevance in pediatric, immunocompromised, and elderly populations. Little is known about hMPV *in vitro* replication processes and their relationship with cellular structures such as the cytoskeleton. Our goal was to evaluate the role of the actin cytoskeleton in hMPV replication at different stages of viral growth. hMPV was isolated in Vero cells from a clinical sample and identified as A_2_ genotype. The cytopathic effect was detected by the appearance of cell rounding and refractory cell clusters. The growth curve showed that viral replication maximum level was between 48 and 72 hpi. The highest percentage of infected cells and intracellular hMPV-protein were detected at the early stages of the replication cycle. Disruption of actin microfilaments with Cytochalasin D during the early events provoked an increase in both intracellular and extracellular viruses. We demonstrate that the early phase of the hMPV curve is crucial for viral replication; and the disruption of microfilaments during this time increments both viral protein expression and release of viruses to the extracellular space. This study contributes to elucidate wild-type hMPV growth kinetics, providing new insights on the actin cytoskeleton role in viral replication mechanisms.

## BACKGROUND

Two members of the *Pneumoviridae* family [1] human Metapneumovirus (hMPV) and respiratory syncytial virus (RSV), cause severe respiratory diseases in infants, immunocompromised and older adults [2]. Currently, for RSV, a bivalent prefusion F vaccine was approved for use during pregnancy in 2023 to protect infants from severe RSV illness during their early months of life [3]. However, there is no licensed vaccine or approved antiviral therapy available for hMPV.

HMPV is an enveloped negative-stranded RNA virus with a non-segmented genome composed of eight genes encoding for nine proteins [4]. Two major hMPV subtypes have been described: A (A1, A2a, A2b1, A2b2) and B (B1, B2) [2,5].

Since the discovery of hMPV [5], several approaches for its *in vitro* isolation have been followed. Viral adaptation and spread in cell cultures are difficult since both require high viral loads and successive blind passages of about 14-21 days to visualize the cytopathic effect (CPE), characterized by changes in cell morphology or by the formation of syncytia [6,7].

The viral infection cycle begins with the entry of particles into susceptible cells through the interaction with receptors that trigger the internalization process. Several studies have reported an important role for the cytoskeleton, particularly actin microfilaments (MFLs) dynamics, in entry and replication of numerous viruses, including hMPV, into the host cell [8]. In this sense, it has been shown that MFLs are essential cofactors for RSV replication, spread, and morphogenesis [9]. Moreover, previous studies from our laboratory demonstrated that the disruption of MFLs during early stage of Pixuna virus (PIXV) replication, increased the extracellular viral yields, probably promoting endocytosis and thus increasing the entry of viral particles [10].

Despite hMPV’s importance as an etiological agent of respiratory pathologies, the mechanisms of interaction between virus and host cell to secure infection remain largely unexplored. We examined the role of MFLs in hMPV replication at different stages of viral growth. To this end, we isolated and identified hMPV circulating in Cordoba, Argentina. Furthermore, we describe the hMPV replication curve and characterize the subcellular expression of viral proteins at different times after infection. We demonstrate that actin depolymerization at the early phase of the hMPV replication curve is crucial for a successful replication. The presence of Cytochalasin D (CytD) during this period increased viral protein expression in the cytoplasm and the release of viruses to the extracellular space.

Considering that hMPV causes a severe disease, identifying the molecular mechanisms underlying its replication cycle broadens our understanding of the cell biology of viral infection. Moreover, it provides possible targets to develop new antiviral treatments.

## METHODS

### Cell culture and Virus isolation

Vero cells (ATCC^®^ CCL-81) were grown in Minimum Essential Medium (MEM, GIBCO-BRL^®^) with 5-10% fetal bovine serum (FBS, Natocor), 1% antibiotic-antimycotic (Pen-Strept 100X, GIBCO), at 37°C and 5% CO_2_.

Human Metapneumovirus was isolated from nasopharyngeal aspirates (NPA) of hospitalized children in Cordoba, Argentina. All procedures were complied with the principles outlined by the Declaration of Helsinki and were approved by an Independent Ethics Committee of Hospital de Niños “Santísima Trinidad” (CIEIS) Protocol: 05/2011. The volunteers who offered samples, signed written assent/consent and their personal data was kept anonymous. The infection protocol was adapted from Van den Hoogen [5]. As infection medium: 0.00125% trypsin (Trypsin Solution 10X, SIGMA^®^), 0.3% bovine serum albumin (BSA, SIGMA^®^) and 1% antibiotic-antimycotic. The NPA positive was diluted in 2 ml of MEM without FBS and centrifuged at 200*Xg* for 5 min (HSR Centrifuge Eppendorf Presvac) (NPA supernatant). Viral inoculum consisted of 150 μl of NPA supernatant plus 50 μl of infection medium. Vero cells (70-80% confluence) were grown in a 24-well plate with MEM. 24 hours later, the plate was washed three times with PBS, 200 μl of the viral inoculum was added in each well and centrifuged at 800*Xg* for 15 min (IEC International Refrigerated Centrifuge Model-PR-2). Afterwards, the plate was incubated at 37°C and 5% CO_2_ for 2 hours, the monolayers were washed three times with PBS and 1 ml of infection medium per well was added. The infection medium was changed at 4, 7, 10, 14, 17, and 21 days post-infection (dpi) and the culture supernatants were stored until processed at -70°C. Cultures were observed daily for CPE by phase-contrast microscopy. Three blind passages for 21 days each were performed. The hMPV positive culture supernatants were used as an infection inoculum.

### RT-PCR

Viral RNA was extracted from 140 μl of culture supernatants using the QIAamp^®^ Viral RNA Mini Kit (Qiagen, GmbH, Hilden, Germany) following the manufacturer’s instructions. RT-PCR, adapted from Bouscambert-Duchamp [11], was carried out as described by Rodriguez [12].

### Indirect immunofluorescence

Infected cells were grown at 60% confluence on glass coverslips during 24 hours. At different times post-infection, cells were washed three times with PBS, fixed with 4% paraformaldehyde and 120 mM sucrose (Sigma-Aldrich Laborchemikalien GmbH, Germany) for 20 min at room temperature (RT). Immunofluorescence (IF) was performed according to Gil [10]. Cells were incubated overnight with mouse anti-F protein/hMPV primary antibody (1/500) (IMAGEN™ hMPV. Oxoid-Ltd.) in 1% BSA/PBS at 4°C. After washed, samples were incubated with goat anti-mouse secondary antibody Alexa-Fluor 488 (1/1600) (Thermo Fisher Scientific Inc., USA) for 1 hour at RT. To visualize the actin filaments and nuclei, cells were labeled with Phalloidin-Tetramethyl rhodamine-B (1/1000) (Sigma-Aldrich) for 1 hour at RT and Hoechst (1X) for 5 min. Then, samples were washed and mounted using Fluorsave (Calbiochem).

### Sequence and phylogenetic analysis

The RT-PCR protocol, adaptaded from Van den Hoogen [13] to amplify a 696 bp fragment of the F gene, was carried out as described by Rodriguez [12]. PCR product was purified with QIAquick Gel Extraction Kit (Qiagen, Germany) following the manufacturer’s instructions. Nucleotide sequencing reactions were performed in both directions using the internal PCR primers by Macrogen, Inc. (Seoul, Korea). Sequences were edited with MEGA-4.0.2 and aligned with sequences available in the GenBank, using the ClustalW. A Maximum Likelihood Tree, based on the amplified region of the fusion protein (F) gene, was generated using PhyML 3.0 software (Université de Montpellier, France). Branch support was evaluated via non-parametric bootstrapping with 1000 pseudoreplicates. The nucleotide substitution model was choosen based on the AIC implemented in ModelTest-3.7 software (University of Vigo, Spain). The sequence was deposited in GenBank (accession no. **MN117139**).

### Viral quantification by RT-qPCR

Viral RNA copies from different supernatants were determined by absolute quantification in an Applied Biosystems 7500 Fast-Real-time PCR system. The reaction mix was prepared by adapting the manufacturer’s protocol for AgPath-ID™ One-Step RT-PCR Reagents (Applied Biosystems™): 2X RT-PCR Buffer= 1X; Forward and reverse PCR primers= 50μM each; Syber Green (SYBR™ Green 10000X-Invitrogen) dilution 1/100= 5.0 and -5; 25X RT-PCR Enzyme Mix (ArrayScript™ Reverse Transcriptase and AmpliTaq Gold^®^DNA Polymerase)= 1X. The final volume was 25 μl, using 2.5 μl of viral RNA. Cycling conditions: 50°C for 30 min; 45 cycles of 94°C for 2 min 95°C for 10 sec, 60°C for 30 sec and 72°C for 30 sec and a final step of 72°C for 10 min. Data were analyzed in real-time using the Virtual Curve qPCR program from Applied Biosystems (Thermo Fisher Connect™). Viral load was calculated from a standard curve generated with serial dilutions (10^−1^ to 10^−12^) of an hMPV N protein synthetic oligonucleotide (199 bp) (Ultramer^®^ DNA Oligo) with a known concentration (6.0219×10^13^ copies of RNA/μl).

### hMPV Replication Curve

Vero cells were grown (60% confluence) on glass coverslips (12mm in 24-well plate) for 24 hours and then infected with isolated hMPV (1.97×10^6^ copies of RNA/μl). The culture supernatants were harvested and the monolayers were fixed at different time points (8, 16, 24, 48, 72, and 96 hpi). Viral RNA was extracted from harvested supernatants, then the extracted RNA was employed for the detection and quantification of hMPV. The mean number of copies of extracellular viral RNA/μl of culture supernatant was then calculated (Applied biosystems. Thermo-Fisher-Connect™).

Cells were processed for IF and the number of fluorescent dots/infected cell were calculated through the analysis of fluorescence images using the ImageJ-Fiji software (ImageJ 1.50c Wayne Rasband. NIH-USA).

### Pharmacological treatment

Vero cells at 60% confluence cultured on glass coverslips were infected with hMPV. Infected cells treated with CytD [2.5μM] [10] or vehicle (infection medium whitout CytD) were analized at different times during the replication cycle. Cells were exposed to CytD or vehicle during the 2 infection hours, as well as throughout the first 8 hpi (experimental period between 0-24 hpi); during the first and the last 24 hours (48-72 hpi) in an experimental period between 0-72 hpi. In each case, the supernatant was removed, the cells were washed three times with PBS and the infection medium was added according to each treatment (Fig. 3B). Culture supernatants and harvested cells were processed for RNA extraction, viral quantification by RT-qPCR and IF, respectively.

### Microscopy and digital image analysis

Immunostained cells were visualized using an inverted epi-fluorescence microscope (Olympus IX81, Olympus). Images were taken with a regular fluorescence microscopy with a CCD camera (Orca 1000, Hamamatsu-Corp.) and with an inverted confocal microscope FV1000 (Olympus). The collection and processing of data from the images were analyzed with the ImageJ-Fiji software. Quantification of intracellular infection was carried out by the percentage of infected cells and the number of fluorescent dots/ infected cell.

### Statistical analysis

All tests and graphs were performed using GraphPad Prism 8.0 (GraphPad Software). Data represent results from at least three independent experiments, and values are presented as the mean ± standard error of the mean (SEM). Assumptions of homoscedasticity and normality were tested using Brown-Forsythe and Barlett’ s tests and D’Agostino & Pearson test, respectively. All assumptions were met, allowing the use of Student’s unpaired t-test or one-way ANOVA test for specific group comparisons. A significance level of p < 0.05 was considered statistically significant.

## RESULTS

### Human-Metapneumovirus isolation and identification

Human Metapneumovirus was isolated from NPA of hospitalized children in Cordoba, Argentina.

The virus replicated in Vero cells cultures after three blind passages of 21 days each one. The CPE was characterized by cell rounding, detachment, and formation of refractory cell clusters (Fig. 1A). IF detection confirmed that the CPE was caused by hMPV. The presence of specific hMPV proteins was observed at 4, 7, and 14 dpi located and distributed in the cell cytoplasm presenting a dotted pattern (Fig. 1B). Simultaneously, virus genome detection by nucleic acid amplification assays (RT-PCR) from supernatants of infected cultures collected at 4, 7, and 14 dpi were positive for the amplification of the 199 bp N protein (Fig. 1C), confirming the presence of hMPV in infected culture. Viral isolation was typified, the Cordoba/ARG/3864/2015 local isolated strain (no: MN117139) was identified as A_2_ hMPV and grouped with Canada and The Netherlands prototype strains, and with sequences from Argentina [12,14], Peru, Brazil, United States, China, and Italy (Fig. 1F).

**Figure 1.**
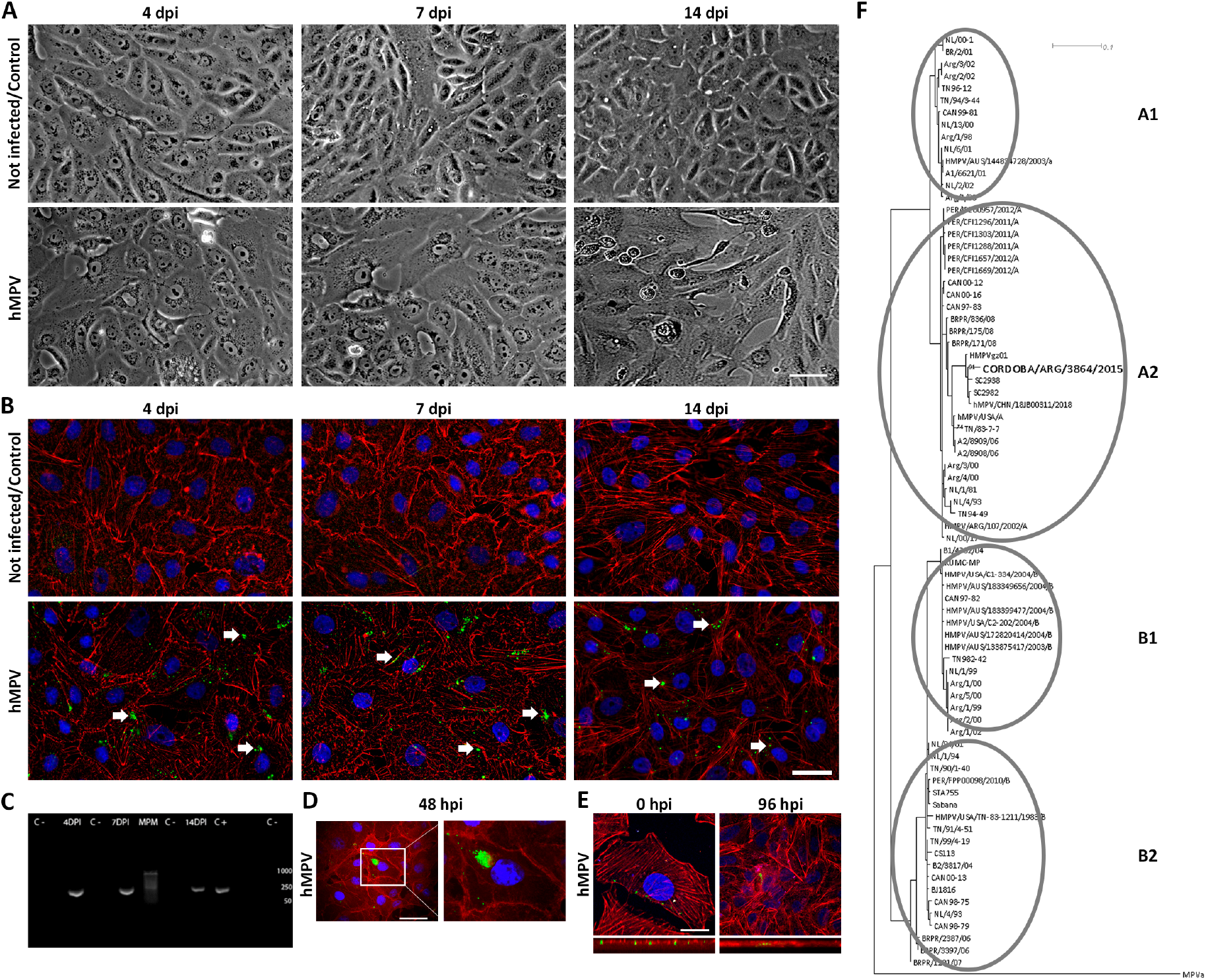
Human Metapneumovirus (hMPV) isolation and identification. A and B: Representative images of uninfected and infected VERO cells with hMPV. **A-** Images were obtained with phase contrast microscopy at low magnification, showing the viral infection cytopathic effect (CPE), which is visible at 14 d.p.i as cell rounding and shedding. Uninfected cells (top panel) and infected cells (bottom panel) at 4, 7 and 14 days dpi. Scale bar: 20 μm. **B-** The localization of the hMPV protein was determined using indirect immunofluorescence with specific anti-hMPV antibodies targeting protein F (green). MFLs were labeled with Phalloidin-Rho (red), and nuclei were stained with Hoechst (blue). A temporary sequence of 4 to 14 days in non infected (top panel) and infected cells (bottom panel). White arrows indicate the fluorescent label of the F protein. Scale bar: 20 μm. **C-** Amplification products (RT-PCR) of the hMPV N protein region (199 bp) by agarose gel electrophoresis. Bands corresponding to culture supernatants of infected cells collected at 4, 7 and 14 dpi and positive controls (C+) are shown. Uninfected cells collected at the same times are included (C-). MPM: molecular weight marker. **D-** Images obtained with epi-fluorescence microscopy, showing that the hMPV F protein accumulates near the nucleus at 48 hpi. Scale bar: 100 μm. **E-** Confocal microscopy images where viral proteins are observed at 0 and 96 hpi. The bottom panel shows a transverse optical section where the viral label is evidenced in the cell cytoplasm decorating the MFLs. Scale bar: 20 μm. **F-** A maximum-likelihood tree (PhyML software) was constructed, using the GTR+G model with parameters suggested by JModelTest 3.7 with bootstraps and 1000 pseudoreplicates. A1: strains to hMPV subgroup A1. A2: strains to hMPV subgroup A2. B1: strains to hMPV subgroup B1. B2: strains to hMPV subgroup B2. Avian MPV (aMPV) was used to root the tree. The strain of interest is marked in bold. The scale bar indicates the changes between nucleotides. All data were obteinded from 5 independet experiments.

Different subcellular localization of large fluorescent dots were observed several times post-infection (Fig. 1B and D), while confocal microscopy detected small dots between 0 and 96 hpi in the subcortical area enriched in actin filaments, below the plasma membrane (Fig. 1E, low panel).

### The hMPV replication cycle

To characterize hMPV replication kinetics, we evaluated the production and release of the virus over time (8, 16, 24, 48, 72, and 96 hpi) using qRT-PCR (RNA copies/µl of culture supernatant). RNA copies increased progressively until 72 hpi, peaking at 4,961 copies/μl of supernatant (Fig. 2A). The time course of hMPV infection was also studied by IF and viral proteins was quantified as number of fluorescent dots/infected cell (Fig. 2B and D). Interestingly, the number of fluorescent dots/infected cell decreased while the number of copies of RNA/μl of culture supernatant increased. At 72 hpi the maximum extracellular viral yield (4,961 copies) corresponded to the minimum number of cytoplasmic dots (2.2 dots), as well as to the lowest percentage of infected cells (32.8%) (Fig. 2C). It is important to highlight that the percentage of infected cells decreased significantly and progressively from 24 hpi until the end of the experiment (Fig. 2C and D).

**Figure 2.**
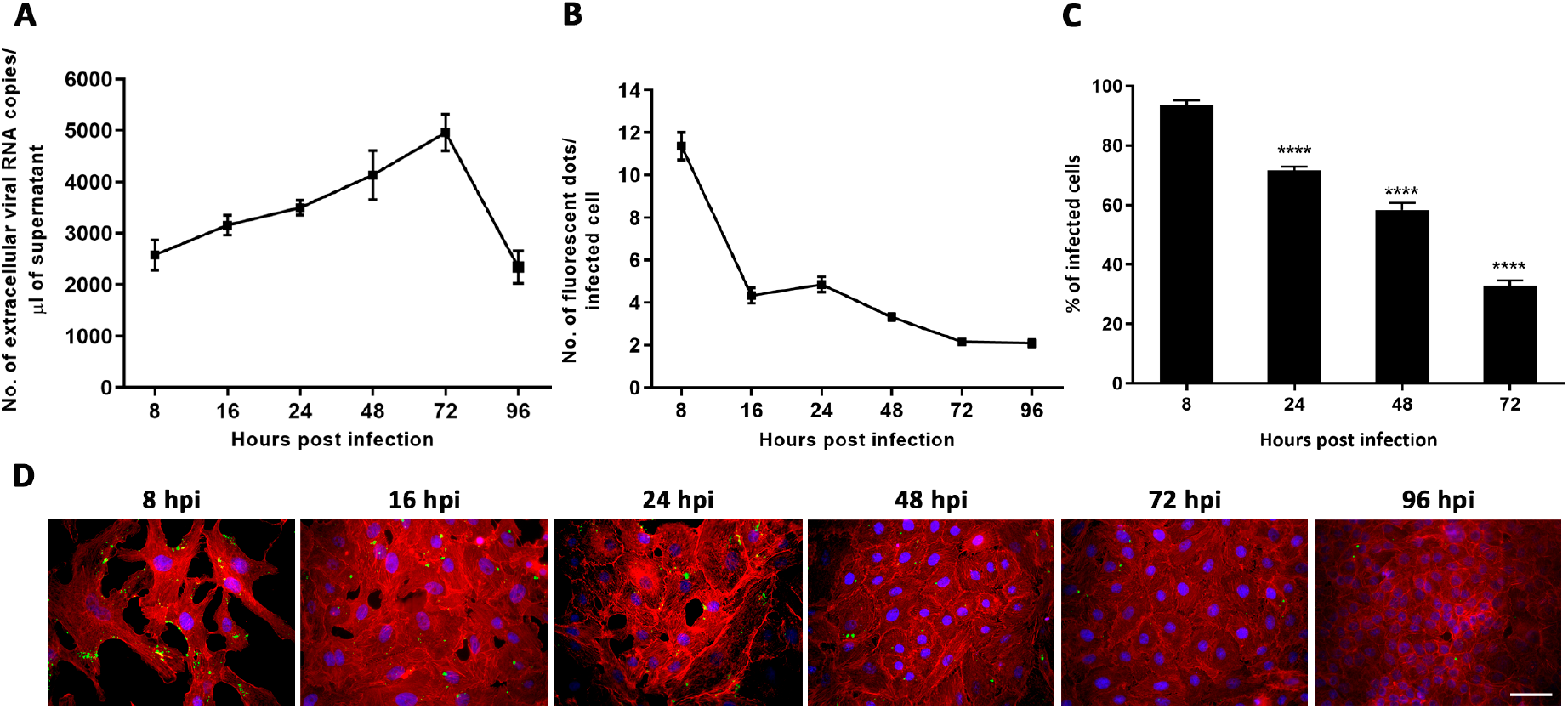
HMPV viral replication curve. **A-** Quantification curve of extracellular viral production through the number of copies of viral RNA in the extracellular medium. Quantification was performed at 8, 16, 24, 48, 72 and 96 hpi. **B-** Quantification of intracellular viral production through the number of fluorescent dots/infected cell (cytoplasmic viral F protein). Quantification was performed at 8, 16, 24, 48, 72 and 96 hpi **C-** Percentage of quantified infected cells was determined by the number of cells with or without fluorescent dots at 8, 24, 48, and 72 hpi. **D-** Representative images of the temporal sequence of infection (0-96 hpi). The hMPV F protein label (green), MFLs (red) and nuclei (Hoechst, blue) are observed. Scale bar: 20 μm. More than 400 cells were screened in each time point. Data represent the mean ± SEM from 3 independent experiments (one-way ANOVA with Tukey test post hoc test). ^****^ <0.0001.

**Figure 3.**
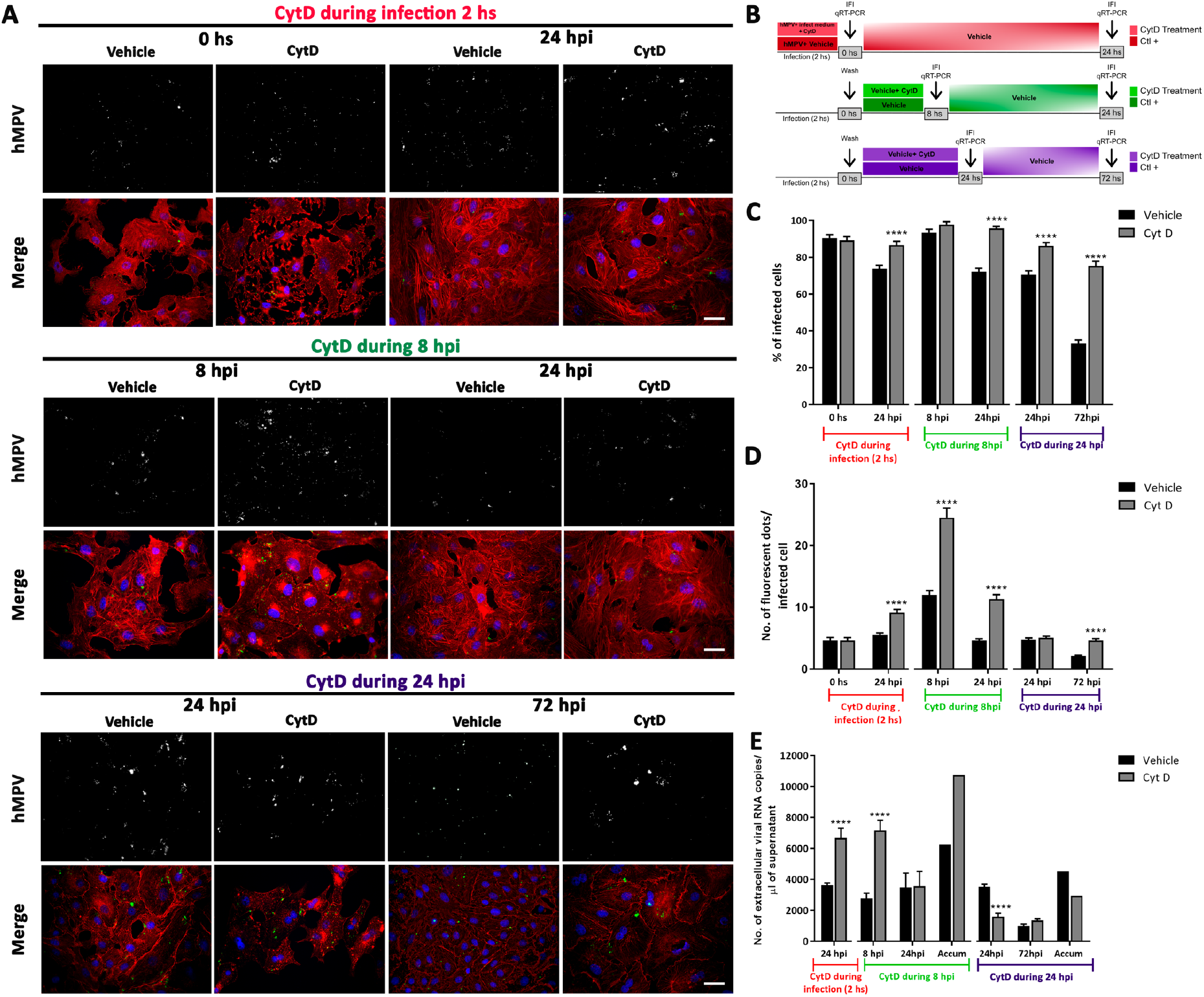
Effect of disruption of MFLs with CytD at different stages of the viral cycle. **A-** Images of CytD treatments in hMPV infected cells at different times of the growth curve: CytD during infection (2 h), during 8 hpi and during 24 hpi. Immunofluorescence performed with specific hMPV antibodies anti-protein F (green), MFLs (Phalloidin-Rho, red) and nuclei (Hoechst, blue). **B-** The scheme illustrates the pharmacological treatment with CytD at different times during the viral replication cycle and the time points of experimental procedures. In red color: cells exposed to vehicle or CytD during the 2 infection hours. In green color: cells exposed to vehicle or CytD throughout the first 8 hpi. In violet color: cells exposed to vehicle or CytD during the first 24 hpi. **C-** Percentage of infected cells after CytD treatment in relation to vehicle control. CytD treatment during infection (2 h): a significant increase was observed at 24 hpi (p< 0.0001). CytD treatment during the first 8 hpi: a significant increase was observed at 24 hpi (p<0.0001). In CytD treatment during first 24 hpi a significant increase was observed both at 24 and 72 hpi (24 hpi: p<0.0001; 72 hpi: p<0.0001). **D-** Number of fluorescent dots per infected cell in CytD treatments and in controls. CytD treatment during infection (2 h): a significant increase was observed at 24 hpi (p< 0.0001). CytD treatment during the first 8 hpi: a significant increase was observed both at 8 and 24 hpi (8 hpi: p< 0.0001; 24 hpi: p< 0.0001). CytD treatment during the first 24 hpi: a significant increase was observed at 72 hpi (p<0.0001). **E-** Quantification of extracellular viral RNA. CytD treatment during infection (2 h): a significant increase in RNA copies in the culture supernatant was observed at 24 hpi (p<0.0001). CytD treatment during the first 8 hpi caused a significant increase in RNA copies in culture supernatant at 8 hpi (p<0.0001); this was also observed in the accumulated values (sum of the means of RNA copies at 8 plus 24 hpi). CytD treatment during the first 24 hpi caused a significant decrease in extracellular viral RNA copies (p< 0.0001), remaining constant until 72 hpi; this was also observed in the accumulated values (sum of the means of RNA copies at 24 and 72 hpi). Scale bar: 20 μm. More than 400 cells were screened in each case. Data represent the mean ± SEM from 3 independent experiments. Student unpaired *t* test was performed (^****^ <0.0001).

Taken together, our results demonstrate that the highest percentage of infected cells and intracellular hMPV-protein contents were detected at the early phase of the replication cycle. While the cycle progressed, hMPV immunostaining decreased whereas extracellular viral RNA increased due to the release of the viral progeny.

### Actin microfilament perturbation at different viral stages affects hMPV replication

Previous studies highlight the involvement of the cytoskeleton, particularly MFLs, in the entry and replication of numerous viruses [8]. Therefore, we studied the role of MFLs at early and late stages of hMPV infection cycle. For this, disruption of actin polymerization was performed using CytD at different times of hMPV infection. We first determined the participation of MFLs during the entry and at its early stages of hMPV replication. To this end, CytD was applied during virus infection (2 hours), during the first 8 hpi and during the first 24 hpi (Fig. 3B).

Treatment with CytD during early replication stages prevented the significant decrease in the number of infected cells, maintaining initial percentage, regardless of drug exposure timing (Fig. 3A and C). Interestingly, CytD treatment during the first 24 hpi, also prevented the dramatic diminution in the percentage of infected cells at 72 hpi (75.3%) compared to control (33.3%) (Fig. 3C). To obtain a quantitative measure of CytD effect on F viral protein expression at the different stages of the replication cycle, we quantified the number of fluorescent dots/infected cell. CytD treatment during infection or during the first 8 hpi significantly increased fluorescent dots/infected cell compared to controls (Fig. 3A and D). Particularly, during the first 8 hpi of treatment a 2 to 2.5 fold increase was observed at both analyzed times (8 hpi and 24 hpi) (Fig. 3A and D). When CytD was applied during the first 24 hpi, and quantified at the end of the treatment, no significant differences in fluorescent dots/infected cells was observed compared to control, but prevented viral protein loss at 72 hpi (Fig. 3D). Finally, we quantified viral-RNA copies in culture supernatant to assess extracellular virus yields (Figure 3E). CytD treatment during the infection significantly increased viral production at 24 hpi. Treatment during the first 8 hpi significantly increased RNA copies at that time. This effect was also observed in the accumulated values, this value is the sum of the mean of RNA copies values at 8 plus 24 hpi (for vehicle: 6,674; CytD treatment: 10,727; see accumulated value at 24 hpi). On the other hand, CytD treatment during the first 24 hpi, caused a significant decrease in extracellular RNA viral copies at 24 hpi, remaining lower up to 72 hpi, as can be observed in the accumulated values (vehicle: 4,510; CytD treatment: 2,930; see accumulated value at 72 hpi) (Fig. 3E).

In addition, we focused on the late stage of the replication curve and observed that CytD applied during the last 24 hpi (48-72 hpi) caused no changes in the percentage of infected cells, in the number of fluorescent dots/infected cell or in the number of extracellular RNA-viral copies compared to controls (data not shown).

These results underscore the importance of the initial 8 hours of hMPV infection for replication, indicating that CytD presence during this period boots viral protein expression in the cell cytoplasm and enhances virus release into extracellular space.

## DISCUSSION

In this study we report the first successful isolation of hMPV performed in Vero cell line from a positive clinical sample in Cordoba, Argentina. We identified this local strain and described the replication curve showing that the maximum viral production takes place at 72 hpi. Moreover, we demonstrate that the disruption of MFL at the early stages of infection increases intracellular viral production and the release of viruses into the extracellular space.

The viral isolate was identified as the A_2_ hMPV subtype through phylogenetic analysis, consistent with our previous studies of A_2_ hMPV subtype circulation in Argentina [12]. Additionally, co-circulation with other genotypes like A_1_, B_1_, and B_2,_ has been reported in our country [14]. Genotype predominance varies based on factors like epidemiological year and region, host immunity, viral load, and susceptibility. Competitive replication was observed between hMPV genotype A and B. A recent study by van den Hoogen’s group [15] demonstrated that lineage A replicates more efficiently than lineage B in an organoid-derived human bronchial-epithelium model, with increased viral-RNA and infectious virus particles. They also highlight the use of viruses with a history of controlled passage, instead of extensively passaged hMPV strains. These data are important because the A_2_ genotype is clinically relevant, as it may cause diseases of varying severity. In studies where verification was possible, genotype A has been associated with severe conditions such as pneumonia and hypoxia, leading to more ICU admissions compared to the other subtypes [16].

In several studies, hMPV replication is limited to certain cell lines, with varied and mild CPE. Isolation of the A_2_ hMPV viral strain in this study showed initial CPE signs at the third blind passage, at 14 dpi, characterized by cell rounding and refractive cells clusters, consistent with previous reports [7,17]. Conversely, other authors have described that hMPV can appear as a syncytium [5,6,18], similar to RSV, and this difference may be related to the cell lines that were infected (e.g. LLC-MK2, tMK, MNT-1, A549) or to the viral genotype [19–21]. Numerous authors note the challenge of hMPV isolation and the absence of an obvious CPE [5,19,20,22]. Isolating respiratory viruses from clinical specimens often exhibits low efficiency *in vitro* due to factors like cell type and virus subtype [17,22–24]. LLC-MK2 and Vero cells are commonly used for all hMPV strains. Although, this study did not compare different cell lines and isolated only the A2 hMPV subtype, efficient viral replication was achieved, consistent with findings by Nao [21] who demonstrated high infectivity with different hMPV strains in VeroE6 and Vero-ATCC cells.

The hMPV replication curve showed an exponential phase between 48 to 72 hpi, with maximum intracellular viral protein concentration at 8 hpi which is consistent with results obtained by other groups. Tollefson et al. [19] described hMPV kinetics in LLC-MK2 cells, with the eclipse phase at 24 hpi and the exponential phase between 48 and 72 hpi with a significant increase in viral titer. El Najjar [25] showed maximum viral production at 72 hpi in BEAS-2B cells, both intra- and extracellularly. Recent studies using recombinant viruses in Vero cells, three-dimensional cultures and organoid-derived bronchial culture have described similar results, both in the CPE caused by the virus and in the replicative cycle [15,16,26]. All these results show that our model of infection in the Vero-CCL cell line with this wild-type virus (A_2_ hMPV), despite its isolation difficulties, is very efficient and representative of *in vivo* infection.

It is known that viruses such as RSV [9] and PIXV [10], among others, take advantage of structures such as the cytoskeleton for entry, replication and exit from the cell. Our study aimed to assess the role of actin cytoskeleton in hMPV replication. For this, we used CytD, known to disrupt actin polymerization. CytD concentration [2.5μM] was effective and consistent with prior studies [24,25] as well as previously in our laboratory with another viral model [10]. The rapid reversible action of CytD facilitated our experimental designs. Intracellularly, CytD not only prevented the decrease in the percentage of hMPV-infected cells (during 2 hours of infection, 8 hpi and 24 hpi) but it also caused a significant increase in the number of fluorescent dots/infected cell, indicative of protein accumulations, in all treatments. In line with this, El Najjar [25] postulates that hMPV proteins assembly during early infection does not require actin polymerization. These protein accumulations (fluorescent dots), resembling “inclusion bodies” [27,28] persist despite actin alteration, hindering viral release. Cifuentes-Muñoz [24] note these structures as primary replication sites, particulary at 24 hpi. In *Pneumovirus*, actin polymerization partially aids inclusion body formation [17]. Unlike Cifuentes-Muñoz [24], our early CytD treatment (8 hpi) boosts extracellular viral load, suggesting that MFLs interruption could facilitate viral entry. This represents an increase in intracellular viral load and a decrease in virus release times, compared to the control. Nevertheless, our results are consistent with those of Cifuentes-Muñoz [24] and El Najjar [25] regarding reduced extracellular viral load from 24 hpi, particularly with continuous 24-hour treatment. Several studies suggest that hMPV relies on cell-cell transmission for efficient spread and infection [8,28–30]. Actin disruption may hinder this process, redirecting virions to alternate exit routes and leading to extracellular accumulation. Therefore, actin dynamics is essential for hMPV infection, enabling cell-to-cell spread regardless of the extracellular viral load [25]. This could explain cytoplasmic viral protein accumulation and the decrease of extracellular viral load after 24 hpi CytD exposure. Given MFLs are involved in viral particle internalization and release [24], it is reasonable to think that short-term disruption of MFLs with CytD would facilitate viral particles internalization. Conversely, prolonged CytD treatment (24 hpi), could prevent the transport of viral proteins and/or nucleocapsid, reducing cytoplasmic accumulation and extracellular viral loading. Despite the years since its discovery, the cell biology of hMPV infection remains poorly understood. This study broadens our knowledge on the isolation and growth curve characterization of a wild-type hMPV obtained from a clinical sample. Furthermore, these findings provide new insights on hMPV infection and the role of actin cytoskeleton in viral replication, enhancing understanding of hMPV internalization and spreading mechanisms.

## ACKNOWLEDGEMENTS

We would like to thank Bernadette van den Hoogen for receiving PER in her laboratory and for advising on the viral isolation protocol; Guadalupe Carballal^†^ and Cristina Videla for generously providing hMPV wild type positive control; Celia Frutos and Lucia Ghietto for technical assistance in Real Time RT-qPCR; Santiago Mirazo for generously providing polyclonal antibodies; Carlos Mas for technical assistance in confocal microscopy at the Centro de Micro y Nanoscopía de Córdoba, CEMINCO-CONICET-Universidad Nacional de Córdoba, Córdoba, Argentina. We particularly want to thank Miss Stella Ascheri for general technical assistance.

## DATA AVAILABILITY STATEMENT

The data that support the findings of this study are available from the corresponding author upon reasonable request.

## REFERENCES

1. Afonso CL, Gaya KA, Bányai K, et al.: TAXONOMY OF THE ORDER MONONEGAVIRALES: UPDATE 2016. Arch Virol 2016; 161:2351–2360.

2. Shafagati N, Williams J: Human metapneumovirus - what we know now. F1000Research 2018; 7:1–11.

3. Kampmann B, Madhi SA, Munjal I, et al.: Bivalent Prefusion F Vaccine in Pregnancy to Prevent RSV Illness in Infants. N Engl J Med 2023; 388:1451–1464.

4. Ballegeer M, Saelens X: Cell-Mediated Responses to Human Metapneumovirus Infection. Viruses 2020; 12:3–22.

5. Van Den Hoogen BG, De Jong JC, Groen J, et al.: A newly discovered human pneumovirus isolated from young children with respiratory tract disease. Nat Med 2001; 7:719–724.

6. Sato K, Watanabe O, Ohmiya S, et al.: Efficient isolation of human metapneumovirus using MNT-1, a human malignant melanoma cell line with early and distinct cytopathic effects. Microbiol Immunol 2017; 61:497–506.

7. Deffrasnes C, Côté S, Boivin G: Analysis of Replication Kinetics of the Human Metapneumovirus in Different Cell Lines by Real-Time PCR. J Clin Microbiol 2005; 43:488–490.

8. Cifuentes-Munoz N, El Najjar F, Dutch RE: Viral cell-to-cell spread: Conventional and non-conventional ways. In: Advances in Virus Research. 2020. p. 85–125.

9. Shahriari S, Wei KJ, Ghildyal R: Respiratory syncytial virus matrix (M) protein interacts with actin in vitro and in cell culture. Viruses 2018; 10:7–12.

10. Gil PI, Albrieu-Llinás G, Mlewski EC, et al.: Pixuna virus modifies host cell cytoskeleton to secure infection. Sci Rep 2017; 7:1–9.

11. Bouscambert-Duchamp M, Lina B, Trompette A, Moret H, Motte J, Andréoletti L: Detection of human metapneumovirus RNA sequences in nasopharyngeal aspirates of young french children with acute bronchiolitis by real-time reverse transcriptase PCR and phylogenetic analysis. J Clin Microbiol 2005; 43:1411–1414.

12. Rodriguez PE, Frutos MC, Adamo MP, et al.: Human Metapneumovirus: Epidemiology and genotype diversity in children and adult patients with respiratory infection in Córdoba, Argentina. PLoS One 2020; 15:e0244093.

13. Van Den Hoogen BG, Osterhaus DME, Fouchier RAM: Clinical impact and diagnosis of human metapneumovirus infection. Pediatr Infect Dis J 2004; 23:25–32.

14. Velez Rueda AJ, Mistchenko AS, Viegas M: Phylogenetic and Phylodynamic Analyses of Human Metapneumovirus in Buenos Aires (Argentina) for a Three-Year Period (2009-2011). PLoS One 2013; 8:1–10.

15. Ribó-Molina P, van Nieuwkoop S, Mykytyn AZ, et al.: Human metapneumovirus infection of organoid-derived human bronchial epithelium represents cell tropism and cytopathology as observed in in vivo models. Virology 2024; 9:1–13.

16. Arnott A, Vong S, Sek M, et al.: Genetic variability of human metapneumovirus amongst an all ages population in Cambodia between 2007 and 2009. Infect Genet Evol 2013; 15:43–52.

17. Kinder JT, Moncman CL, Barrett C, Jin H, Kallewaard N, Dutch RE: Respiratory Syncytial Virus and Human Metapneumovirus Infections in Three-Dimensional Human Airway Tissues Expose an Interesting Dichotomy in Viral Replication, Spread, and Inhibition by Neutralizing Antibodies. J Virol 2020; 94:1–20.

18. Bernal LJ, Velandia-Romero M, Guevara C, Castellanos JE: Human Metapneumovirus: Laboratory Methods for Isolation, Propagation, and Plaque Titration. Intervirology 2019; 61:301–306.

19. Tollefson SJ, Cox RG, Williams J V: Studies of culture conditions and environmental stability of human metapneumovirus. Virus Res 2010; 151:54–59.

20. Jumat MR, Nguyen Huong T, Wong P, et al.: Imaging analysis of human metapneumovirus-infected cells provides evidence for the involvement of F-actin and the raft-lipid microdomains in virus morphogenesis. Virol J 2014; 11:1–12.

21. Nao N, Sato K, Yamagishi J, et al.: Consensus and variations in cell line specificity among human metapneumovirus strains. PLoS One 2019; 14:1–21.

22. Lee H, Woo HM, Kim K, et al.: Improving pneumovirus isolation using a centrifugation and AZD1480 combined method. J Microbiol Biotechnol 2019; 29:2006–2013.

23. Cox RG, Mainou BA, Johnson M, et al.: Human Metapneumovirus Is Capable of Entering Cells by Fusion with Endosomal Membranes. PLoS Pathog 2015; 11:1–29.

24. Cifuentes-muñoz N, Branttie J, Slaughter KB, Dutch RE: Human Metapneumovirus Induces Formation of Inclusion Bodies for Efficient Genome Replication and Transcription. J Virol 2017; 91:1–18.

25. El Najjar F, Cifuentes-Muñoz N, Chen J, et al.: Human metapneumovirus Induces Reorganization of the Actin Cytoskeleton for Direct Cell-to-Cell Spread. PLoS Pathog 2016; 12:1–30.

26. Geiser J, Boivin G, Huang S, et al.: RSV and HMPV Infections in 3D Tissue Cultures: Mechanisms Involved in Virus-Host and Virus-Virus Interactions. Viruses 2021; 13:1–15.

27. Derdowski A, Peters TR, Glover N, et al.: Human metapneumovirus nucleoprotein and phosphoprotein interact and provide the minimal requirements for inclusion body formation. J Gen Virol 2008; 89:2698–2708.

28. Najjar F El, Castillo SR, Moncman CL, et al.: Imaging analysis reveals budding of filamentous human metapneumovirus virions and direct transfer of inclusion bodies through intercellular extensions. MBio 2023; 14:1–18.

29. Merwaiss F, Czibener C, Alvarez DE: Cell-to-cell transmission is the main mechanism supporting bovine viral diarrhea virus spread in cell culture. J Virol 2018; 93:1–18.

30. Boggs KB, Edmonds K, Cifuentes-Munoz N, et al.: Human Metapneumovirus Phosphoprotein Independently Drives Phase Separation and Recruits Nucleoprotein to Liquid-Like Bodies. MBio 2022; 13:.

